# Novel Insights into Malaria Vector Surveillance in Madagascar Using a Quadrant Enabled Screen Trap (QUEST) and Bloodmeal Detection Assay for Regional Transmission (BLOODART)

**DOI:** 10.1101/530758

**Authors:** Riley E. Tedrow, Tovonahary Rakotomanga, Thiery Nepomichene, Jocelyn Ratovonjato, Arséne C. Ratsimbasoa, Gavin J. Svenson, Peter A. Zimmerman

## Abstract

**Background:** The Madagascar National Strategic Plan for Malaria Control 2018 (NSP) outlines malaria control pre-elimination strategies that include detailed goals for mosquito control. Primary surveillance protocols and mosquito control interventions focus on indoor vectors of malaria, while many potential vectors feed and rest outdoors. Here we describe the application of novel tools that advance our understanding of diversity, host choice, and *Plasmodium* infection in the Anopheline mosquitoes of the Western Highland Fringe of Madagascar.

**Methodology/Principal Findings:** We employed a novel outdoor trap, the QUadrant Enabled Screen Trap (QUEST), in conjunction with the recently developed multiplex BLOOdmeal Detection Assay for Regional Transmission (BLOODART). We captured a total of 1252 female *Anopheles* mosquitoes (10 species), all of which were subjected to BLOODART analysis. QUEST collection captured a heterogenous distribution of mosquito density, diversity, host choice, and *Plasmodium* infection. Concordance between *Anopheles* morphology and BLOODART species identifications ranged from 93-99%. Mosquito feeding behavior in this collection frequently exhibited multiple blood meal hosts (single host = 53.6%, two hosts = 42.1%, three hosts = 4.3%). The overall percentage of human positive bloodmeals increased between the December 2017 and the April 2018 timepoints (27% to 44%). *Plasmodium* positivity was found primarily in vectors considered to be of secondary importance, with an overall prevalence of 6%.

**Conclusions/Significance:** The QUEST was an efficient tool for sampling Anopheline mosquitoes. Vectors considered to be of secondary importance were commonly found with *Plasmodium* DNA in their abdomens, indicating a need to account for these species in routine surveillance efforts. Mosquitoes exhibited multiple blood feeding behavior within a gonotrophic cycle, with predominantly non-human hosts in the bloodmeal. Taken together, this complex feeding behavior could enhance the role of multiple Anopheline species in malaria transmission, possibly tempered by zoophilic feeding tendencies.

**Author Summary:** Malaria continues to be a significant threat to public health in Madagascar. Elimination of this disease is impeded by numerous factors, such as vector surveillance that does little to account for the potential role of secondary malaria vectors, which rest and feed outdoors. In this study, we designed a novel, low cost QUadrant Enabled Screen Trap (QUEST) to address the lack of traps for outdoor mosquitoes. We used this in conjunction with our novel BLOOdmeal Detection Assay for Regional Transmission (BLOODART) to assess mosquito feeding behavior in the Western Highland Fringe of Madagascar. Our analysis revealed significant variability in mosquito density, diversity, host choice, and *Plasmodium* infection across traps placed within and between two nearby villages at two timepoints; indicating a strong, small-scale spatial component to disease transmission that warrants further investigation. Many of the mosquitoes in this sample (46.4%) fed on two or three host species, indicating complex feeding behaviors that could influence malaria transmission. Further, *Plasmodium* DNA was detected in the abdomens of numerous vectors of supposed secondary importance, indicating a neglected parasite reservoir and an increased need to account for these species in routine surveillance efforts.

## Introduction

The Madagascar National Strategic Plan for Malaria Control 2018 (NSP), developed in coordination with the Madagascar National Malaria Control Program (NMCP), outlines pre-elimination strategies and a plan of action for malaria control in Madagascar [1]. The priorities of this plan reflect lessons learned from a century of malaria control efforts in the country. Between World Wars I and II, the antimalaria service of Madagascar implemented 1.) limited prophylaxis, 2.) mosquito larval control with Paris Green insecticide, 3.) the introduction of mosquito larvae-eating fish (*Gambusia* sp.) to cisterns and irrigation ditches, and 4.) drainage works to limit mosquito breeding sites [2,3]. The first major success followed the 1950s introduction of the insecticide DDT to the Central Highlands for indoor residual spraying [4]. This campaign combined DDT and chemoprophylaxis to achieve interruption of transmission in the region [5]. Much of this success may be attributed to the elimination of the primary vector (*Anopheles funestus*) from the highlands by 1952 [6]. Believing the intervention complete, spraying ceased in 1975. The following decade was characterized by the rapid deterioration of the health network through the erosion of health facilities, drug stock outages, and medical staff absenteeism [7–10]. A plea for the reintroduction of control measures followed the 1986 discovery of a single *Plasmodium* positive *An. funestus* in the highlands [9]. No action was taken, and an epidemic followed that took the lives of ~40,000 people [5,11,12].

Today, an evolved perspective of the entomological side of the 1980s epidemic has emerged. Spraying campaigns are never truly comprehensive, leaving reservoirs that facilitate future recolonization events [12]. *Anopheles funestus* populations from the highlands are genetically similar to populations on the east coast [13], suggesting that the highlands reinvasion of *An. funestus* may have come from the east. Furthermore, it is likely that additional Anopheline vector species contributed to the malaria outbreak [9]. Mosquitoes are often classified as primary, secondary, and accidental vectors; assigned by purported importance for malaria transmission in a particular region [14,15]. These loosely-defined designations likely contribute to a knowledge gap between primary and secondary vectors.

Numerous secondary vectors, present in Madagascar, have been documented with *Plasmodium* sporozoites by dissection of the salivary glands: An. *coustani* [14,16], An. *mascarensis* [17], An. *squamosus* [16,18], An. *pharoensis* [16], An. *rufipes* [14], and An. *maculipalpis* [14]. Circumsporozoite positive ELISA further implicates *An. coustani* [10,19,20], *An. mascarensis* [10,17,21], and *An. squamosus* [22] as malaria vectors. While common sampling strategies are biased toward endophilic and endophagic mosquitoes [23], most of these potential Malagasy malaria vectors are both exophagic and exophilic [6,21,24]. Detailed goals of the NSP describe objectives for entomological surveillance of sentinel sites in Madagascar, seeking data on vector taxonomy, density, feeding behavior, insecticide susceptibility, parity, age, and sporozoite rate. Here we seek to advance our understanding of the diversity, feeding behavior, and *Plasmodium* infection status of potential malaria vectors in the Western Highland Fringe of Madagascar to achieve a more comprehensive picture of malaria transmission in the region. To accomplish this goal, we utilized the recently developed Bloodmeal Detection Assay for Regional Transmission (BLOODART), designed to simultaneously assess *Anopheles* mosquito species, mammalian hosts, and *Plasmodium* parasites from an excised mosquito abdomen or abdomen squash [25].

In conjunction with BLOODART, we employed a novel barrier screen trap, the Quadrant Enabled Screen Trap (QUEST). Standard barrier screens represent a relatively recent collection method, designed to address the paucity of effective and unbiased collection tools for exophilic mosquitoes [26]. Outdoor traps typically attempt to replicate existing outdoor resting spots for mosquitoes, requiring them to compete with potentially more abundant natural options [27,28]. Further, seeking resting mosquitoes by manually searching amongst the vegetation requires significant time and effort for a small return on samples [29,30]. Standard barrier screens offer the advantage of being permeable to visual and olfactory cues, perhaps making them less likely to divert the mosquito [26]. These screens appear to intercept mosquitoes irrespective of species-specific resting site or host preferences [31], reducing the potential for bias. This may be reflected in the equal or greater Anopheline diversity captured on barrier screens compared to human landing catches [26]. Standard barrier screens provide some sense of directionality by assessing which side of the net mosquitoes were captured from. We sought to provide greater directional resolution and capture numbers than a standard barrier screen by designing a cross-shaped barrier screen with built in eaves. As mosquitoes tend to move up and over physical barriers [32], we suspect the addition of eaves might prolong mosquito detainment. By superimposing our QUEST captured mosquitoes and BLOODART analysis, this study aims to assess mosquito density, diversity, host choice, and *Plasmodium* positivity with spatial resolution, demonstrating the ability of these tools to contribute to the entomological surveillance goals of the NSP [1].

## Methods

### Mosquito Field Collection

Wild-caught mosquitoes were collected by Case Western Reserve University and Madagascar NMCP entomologists in December 2017 (six nights) and April 2018 (five nights), corresponding to either side of the rainy season (December to March [33]). A peak in clinical malaria has been observed in April-May for the Tsiromandidy Health District [34]. Collections were performed in the villages of Amparihy and Ambolodina, located in the fokontany of Kambatsoa (Commune Maroharona, Tsiroanomandidy Health District). Epidemiological surveys have been previously performed in this area in partnership with the Madagascar NMCP [35], and are consistent with protocols approved by the Madagascar Ministry of Health for the present study (N°099-MSANP/CE). Additionally, community and household approvals were obtained following fokontany-based meetings prior to initiating all study activities.

Mosquitoes were collected using QUEST, modified from a previously described barrier screen design [26]. One round of indoor pyrethroid spray catch was conducted in Amparihy (10 dwellings) and Ambolodina (seven dwellings) in December 2017 as previously described [25], with permission from the owner of the residence. QUEST collections began at 18:00 hrs and continued at three-hour increments with a final collection at 06:00 hrs the following day, for a total of five sampling events. Mosquitoes were aspirated on a quadrant-by-quadrant basis and deposited live into pre-labeled paper cups at the beginning of each collection timepoint. Collected mosquitoes were incapacitated by ether and keyed to species using local unpublished keys from Institut Pasteur de Madagascar. Mosquitoes were preserved in individual 2 ml tubes containing 90% ethanol with a label including a unique specimen voucher code containing the prefix “APY.” A subset of mosquitoes in April 2018 were collected on filter paper (n=115) after ethanol supplies were exhausted.

### Trap Design and Placement

QUESTs were designed to improve yields and increase our understanding of mosquito mobility within the study site (Fig 1). Poles for erecting the traps were sourced from trees onsite. The net material used for the traps was a flexible white fabric similar to an untreated bed net. Traps were cross-shaped (aerial view), with each extension measuring 5 m in length and approximately 1.5 m tall. The top of each extension was affixed with a 0.25 m overhang tied at a 45° angle, creating an “eave” to trap insects attempting to bypass the barrier by flying up and over. The design provided four isolated quadrants, with each extension pointing in a cardinal direction. The quadrants were uniformly numbered as follows: 1-Northwest, 2-Northeast, 3-Southwest, 4-Southeast. Three QUEST were set up in a village (Amparihy Trap 1 – Trap 3, Ambolodina Trap 1 – Trap 3), in close proximity to homes and livestock. A single standard barrier screen [26] (Ambolodina Trap 4; Fig 1E,1F) was installed inside a cattle pen in the December 2017 collection. This trap was approximately 1.5 m tall and 15 m in length, built with locally sourced poles and a similar net-like material. We used three QUESTs for 11 nights, therefore a total of 33 trap-nights. The standard barrier screen was used for two nights.

**Fig 1.**
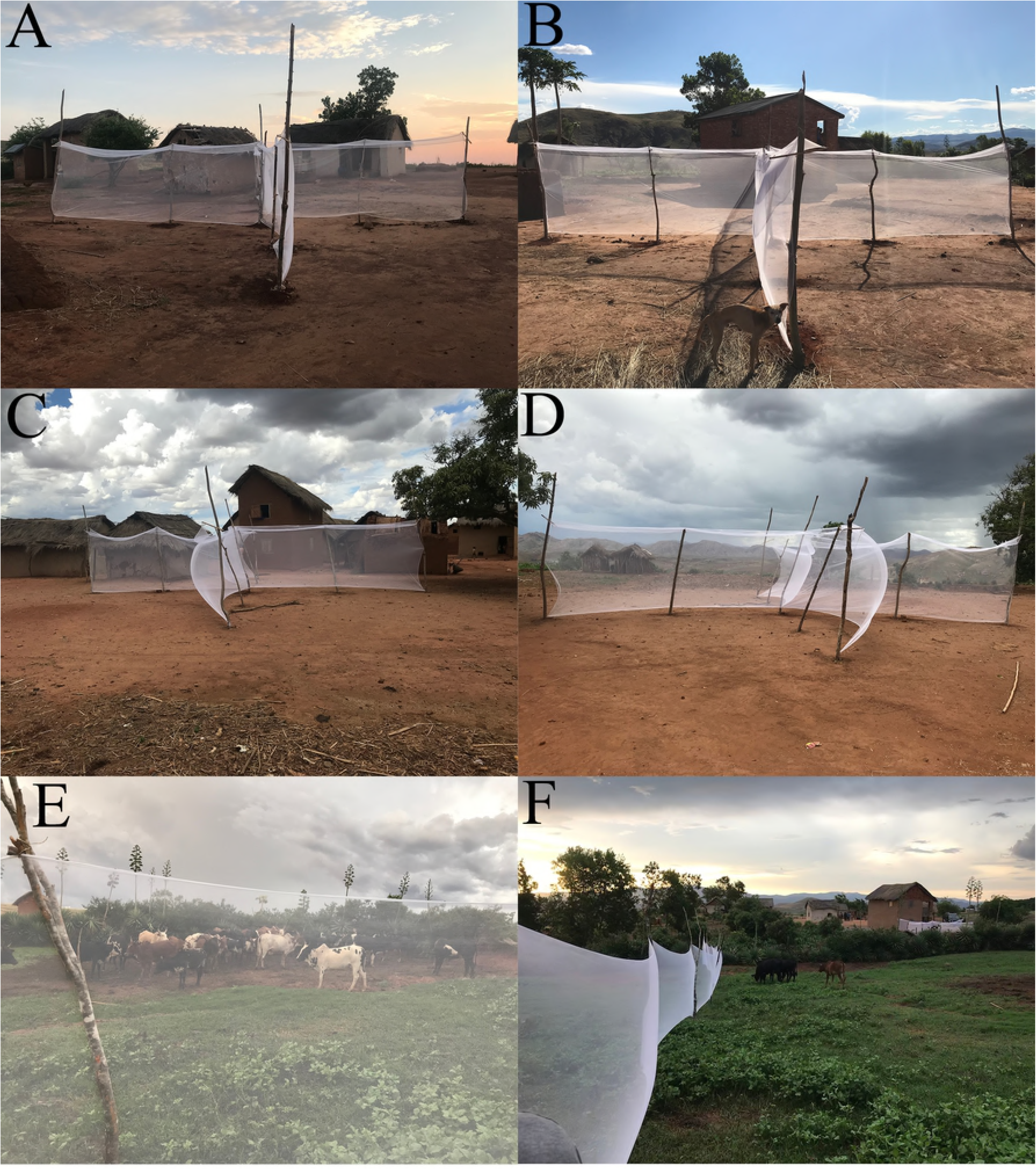
Quadrant Enabled Screen Trap (QUEST) with built-in eaves. A,B: Ambolodina T1. C, D: Amparihy T1. Standard Barrier Screen. E,F: Ambolodina T4.

### Trap Coordinates

QUEST (Quadrant Enabled Barrier Screen) (Fig 1A-D)

Amparihy Trap 1: S 19°23.112’, E 046°08.186’; 763m altitude

Amparihy Trap 2: S 19°23.089’, E 046°08.176’; 775m altitude

Amparihy Trap 3: S 19°23.059’, E 046°08.220’; 779m altitude

Ambolodina Trap 1: S 19°22.460’, E 046°07.459’; 808m altitude

Ambolodina Trap 2: S 19°22.457’, E 046°07.405’; 803m altitude

Ambolodina Trap 3: S 19°22.458’, E 046°07.356’; 802m altitude

Standard Barrier Screen (Fig 1E-F)

Ambolodina Trap 4: S 19°22.466’, E 046°07.439’; 809m altitude

### Mosquito Photography

High-resolution images of preserved specimens were captured using a Passport Storm© system (Visionary DigitalTM, 2012), including: a Stackshot z-stepper, a Canon 5D SLR, a MP-E 65 mm macro lens, three Speedlight 580EX II flash units, and a computer running Canon utility and Adobe Lightroom 3.6 software. The z-stepper was controlled through Zerene Stacker 1.04 and images were stacked using the P-Max protocol. To prepare for photography, the specimen was removed from EtOH, allowed to dry for several minutes, and temporarily affixed to a pin with a small dab of K-Y™ Personal Lubricant. Images were captured over an 18% gray card background and processed in Adobe Photoshop CC 2018 to adjust lighting and sharpness, and to add scale bars. Minor adjustments were made using the stamp tool to correct background and stacking aberrations.

### Extraction and PCR

DNA extraction and PCR amplification of targets for mosquito species, mammalian host, and *Plasmodium* parasites were carried out as previously described [25].

### BLOODART

We utilized the Bloodmeal Detection Assay for Regional Transmission [25] with the addition of several new probes, including *An. squamosus*, two probes that distinguish *An. gambiae sensu stricto* from *An. arabiensis*, ringtail lemur (*Lemur catta*), Coquerel’s sifaka (*Propithecus coquereli*), and the house mouse (*Mus musculus*). A list of probes and fluorescent microspheres for the mammalian and mosquito probes is located in S1 Table. Probes for *Plasmodium* species are described elsewhere [36].

### Proboscis measurement

To determine the effect of proboscis length on host choice, we measured the proboscis and wing length of 117 *An. coustani*, 58 of which were exclusively human-blood positive, and 59 that were exclusively cow-blood positive. Measurements followed the protocol of Wheeler, 1993 [37], with wing length used as a proxy for body size.

### Data analysis

Statistical analyses, histograms, and scatterplots were generated in R Version 3.4.0 [38] using the compilation package ‘Tidyverse’ [39]. Data analyzed in this study has been added to VectorBase (VBP0000345).

## Results

### Mosquito Capture

We captured 501 mosquitoes in December 2017, and 751 mosquitoes in April 2018. The QUEST captured 1143 mosquitoes over 33 trap-nights (15 Amparihy, 18 Ambolodina) for 34.6 mosquitoes per trap/night. Overall, there was a greater concentration of mosquitoes on the traps at the April 2018 timepoint. The distribution of mosquitoes within each trap differed, with some QUESTs showing a homogenous distribution of mosquitoes across the four quadrants while others had a clear concentration of samples on a subset of the four quadrants (Fig 2, S2 Table). The standard barrier screen captured 101 mosquitoes in two nights, thus a total of 50.5 mosquitoes per trap/night. On the standard barrier screen, 91% of mosquitoes were captured on the same side of the net where the Malagasy zebu rested and slept. The standard barrier screen captured seven of the 10 Anopheline species observed on QUEST (S3 Table). However, the species missing for this trap had low overall sample numbers. Mosquitoes began to appear on the QUESTs at the 21:00 hrs collection point (29.3%), peaked at 00:00 hrs (32.9%), and tapered off into the early morning collections at 03:00 hrs (25.5%) and 06:00 hrs (12.3%). The mosquito activity pattern for individual nights was variable. No mosquitoes were captured at 18:00 hrs. Only eight mosquitoes were captured by pyrethroid spray catch across 17 dwellings, with all mosquitoes identified as *An. funestus*. We have elected to combine mosquitoes from all three capture methods for subsequent analyses, with results from specific traps designated as such in the text.

**Fig 2.**
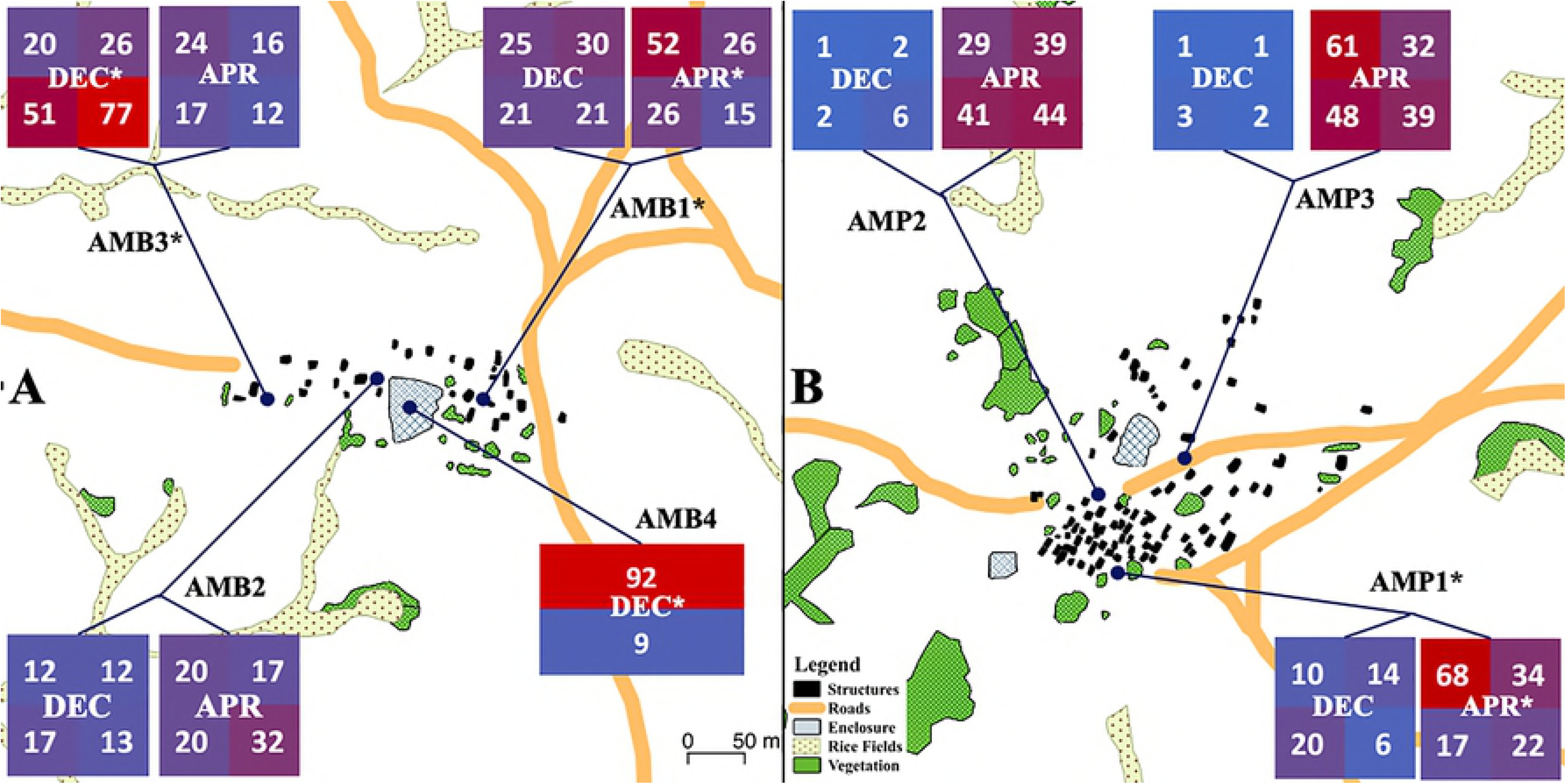
Map of villages with trap placement. Each heatmap box represents the count of mosquitoes distributed across the four quadrants of the trap for December 2017 or April 2018. AMB1-3: Ambolodina (AMB) Trap 1 – Trap 3 QUEST. AMB4: Ambolodina Trap 4 standard barrier screen. Time points marked with an asterisk indicate a non-random distribution of mosquitoes across the trap at that timepoint. Traps marked with an asterisk indicate significantly different mosquito distributions when comparing the two timepoints for the same trap. AMB4 laterally bisected the cattle enclosure it was placed in. AMP1-3: Amparihy (AMP) Trap 1 – Trap 3 QUEST. Map generated using QGIS v.2.18. Numerical version available in S2 Table.

### Mosquito Characterization

The mosquito probe set used in this study was designed to capture species detected in prior surveys conducted in our study villages [25]. As they cannot be morphologically distinguished, *An. gambiae s.s*. and *An. arabiensis* were considered as the *An. gambiae sensu lato* complex in our comparison of morphological versus BLOODART identifications. Further data analyses here preferentially used the species determination of BLOODART over morphology. Where BLOODART identifications were inconclusive, we used the original morphological identification. Of the 64 mosquitoes designated as “inconclusive” by BLOODART, 16 produced a clean PCR band, 24 produced either a smear or several bands of unexpected size, five produced faint bands, and 19 produced no band at all. Concordance for individual probes ranged from 93-99% (Table 1). The unknown species probe of Tedrow, 2018 [25], hybridized primarily to specimens morphologically identified as *An. mascarensis* (22/34), and will be considered in this manuscript as the *An. mascarensis* probe. We provide several photographs (Figs 3 and 4) of Anopheline species captured in this study (prior to processing for BLOODART analysis). This provides a morphological representative for the molecular probes used in BLOODART, and furthermore, provides publicly available images for several of these poorly studied species.

**Table 1.** Counts of mosquitoes identified by morphology and BLOODART.

| Morphology | BLOODART |  |  |  |  |  |  |  | Total |
| --- | --- | --- | --- | --- | --- | --- | --- | --- | --- |
|  | *coustani* | *maculipalpis* | *squamosus* | *rufipes* | *gambiae s.l.* | *mascarensis* | *funestus* | Inconclusive |  |
| *coustani* | **433** | 14 | 6 | 3 | 3 | 1 | 1 | 4 | 465 |
| *maculipalpis* | 1 | **190** | 9 | 1 | 0 | 1 | 1 | 14 | 42 |
| *squamosus* | 12 | 9 | **172** | 4 | 1 | 2 | 0 | 35 | 78 |
| *rufipes* | 9 | 30 | 1 | **123** | 3 | 1 | 2 | 4 | 217 |
| *gambiae s.l.* | 3 | 0 | 2 | 1 | **70** | 1 | 1 | 0 | 173 |
| *mascarensis* | 2 | 3 | 0 | 6 | 0 | **22** | 0 | 1 | 34 |
| *funestus* | 4 | 2 | 3 | 5 | 1 | 6 | **20** | 1 | 235 |
| *flavicosta* | 0 | 0 | 0 | 0 | 0 | 0 | **0** | 1 | 1 |
| *fuscicolor* | 0 | 1 | 0 | 0 | 0 | 0 | **0** | 0 | 1 |
| *pretoriensis* | 1 | 1 | 0 | 0 | 0 | 0 | **0** | 4 | 6 |
| TOTAL | 465 | 250 | 193 | 143 | 78 | 34 | 25 | 64 | 1252 |
| Sensitivity | 93% | 76% | 89% | 86% | 90% | 65% | 80% |  |  |
| Specificity | 96% | 97% | 94% | 95% | 99% | 99% | 98% |  |  |
| Concordance | 95% | 93% | 93% | 94% | 99% | 98% | 98% |  |  |

**Fig 3.**
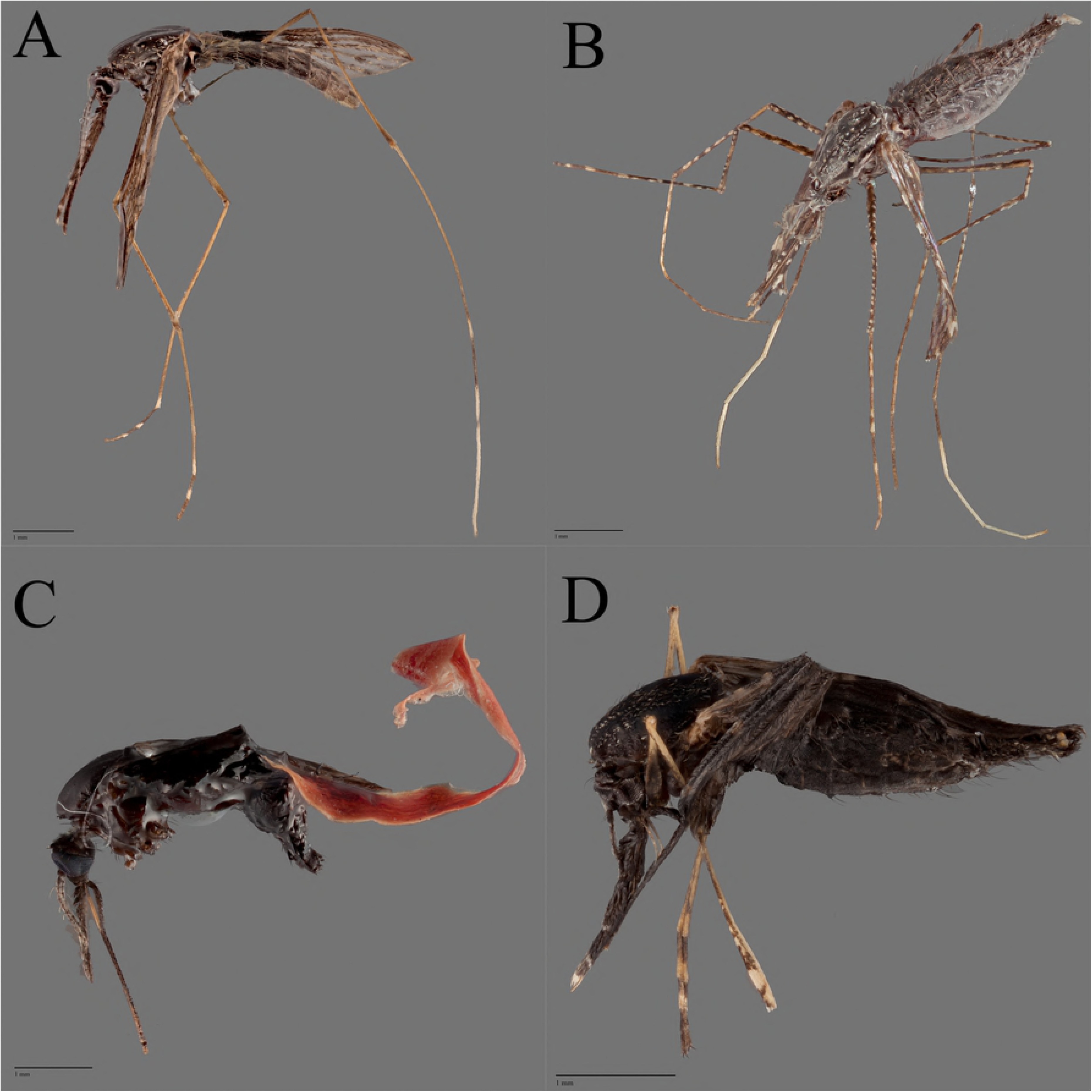
High-resolution z-stack image of ethanol-stored mosquito specimens 1. A: *Anopheles coustani* (APY1071), B: *Anopheles maculipalpis* (APY684), C: *Anopheles rufipes* (APY1016), D: *Anopheles squamosus* (APY702).

**Fig 4.**
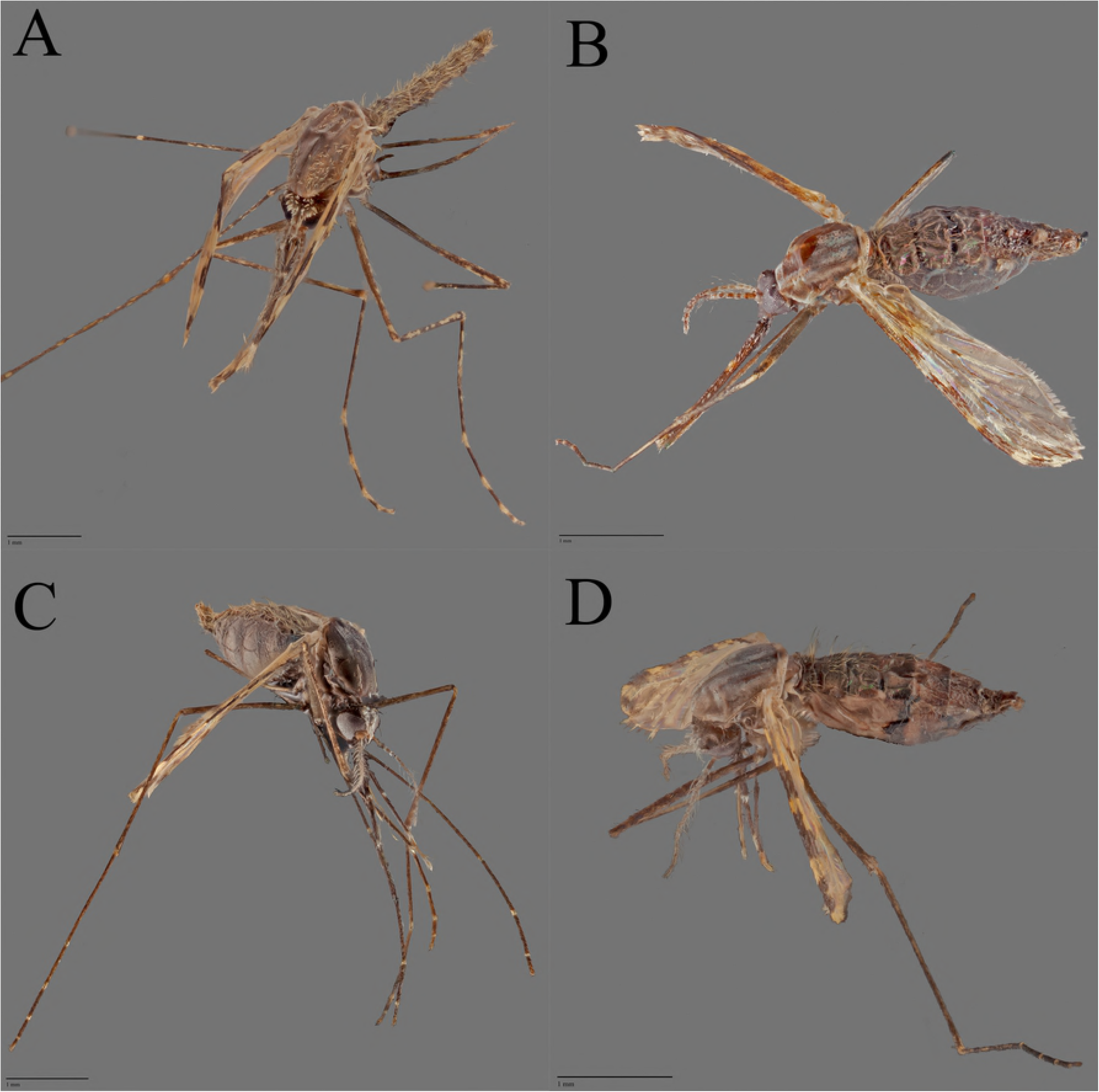
High-resolution z-stack image of ethanol-stored mosquito specimens 2. A: *Anopheles arabiensis* (APY689), B: *Anopheles funestus* (APY237), C: *Anopheles mascarensis* (APY437), D: *Anopheles mascarensis* (APY733).

### Mosquito Abundance and Diversity

The predominant mosquito species, pooling all trapping methods, were *An. coustani* (n=469, Fig 3A), *An. maculipalpis* (n=263, Fig 3B), *An. rufipes* (n=147, Fig 3C), and *An. squamosus* (n=228, Fig 3D). Species captured in lower numbers were *An. arabiensis* (n= 64, Fig 4A), *An. mascarensis* (n=35, Fig 4C, D), *An. funestus* (n=26, Fig 4B), *An. gambiae s.s*. (n=14), *An. pretoriensis* (n=5), and *An. flavicosta* (n=1). The abundance of particular species shifted on either side of the rainy season, with an even mix of *An. arabiensis* and *An. gambiae s.s*. in December 2017 shifting to exclusively *An. arabiensis* in April 2018. The proportion (Fig 5, S4 Table) and distribution (Fig 2, S2 Table) of *Anopheles* species on the QUESTs was complex. There were nearly seven times more mosquitoes captured in Amparihy in April 2018 compared to December 2017. In April 2018, traps in Amparihy collected proportionally more *An. arabiensis*, *An. coustani*, and *An. funestus* than the traps in Ambolodina, while the proportion of *An. rufipes* declined.

**Fig 5.**
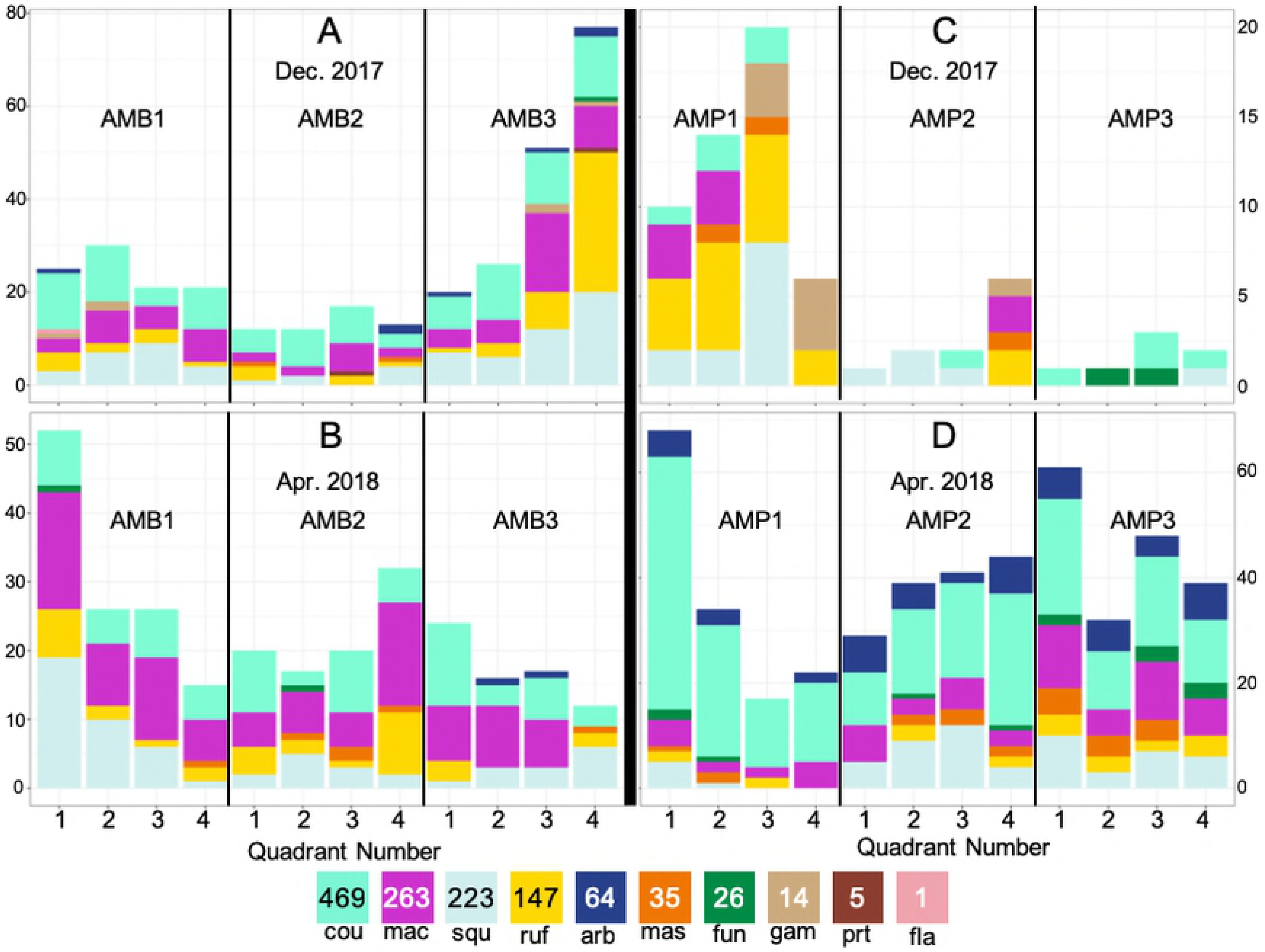
Distribution of mosquito species across QUEST. Three-letter combinations indicate mosquito species: arb =*An. arabiensis*, cou =*An. coustani*, fun =*An. funestus*, fla =*An. flavicosta*, gam =*An. gambiae*, mac =*An. maculipalpis*, mas =*An. mascarensis*, prt =*An. pretoriensis*, ruf =*An. rufipes*, squ =*An. squamosus*. Quadrant numbers refer to the direction of their orientation: 1-Northwest, 2-Northeast, 3-Southwest, 4-Southeast. (A) Ambolodina December 2017, (B) Ambolodina April 2018, (C) Amparihy December 2017, (D) Amparihy April 2018. Numerical version available in S4 Table.

### Host Choice

Mosquitoes in this collection exhibited complex blood feeding behaviors. Six mammalian hosts were detected across our blood meal samples, with the predominant constituents coming from bovine (90.3%), followed by human (39.5%) and porcine (15.3%) blood (Fig 6). Canine (4.7%), feline (0.3%) and hircine (0.1%) blood were detected infrequently. No mosquito was positive for lemur or murine blood. In addition to evidence of many blood meal hosts, individual mosquitoes frequently harbored blood from two or three different host species: single host = 53.6%, two hosts = 42.1%, three hosts = 4.3%. The incidence of multiple blood feeding was significantly different (χ^2^= 42.3, 2 DOF, p < 0.0001) from December 2017 (single host = 41.8%, two hosts = 52.3%, three hosts = 5.8%) to April 2018 (single host = 60.9%, two hosts = 35.7%, three hosts = 3.2%). The most anthropophilic species were *An. coustani* and *An. mascarensis*. The propensity for human feeding increased significantly from the December 2017 (27% human positive) to the April 2018 (44% human positive) collection (χ^2^= 44.8, 1 DOF, p < 0.0001), with anthropophilic shifts observed for *An. coustani* (32% to 60%), *An. funestus* (9% to 50%), *An. rufipes* (23% to 44%), and *An. arabiensis* (38% to 53%) (Fig 6).

**Fig 6.**
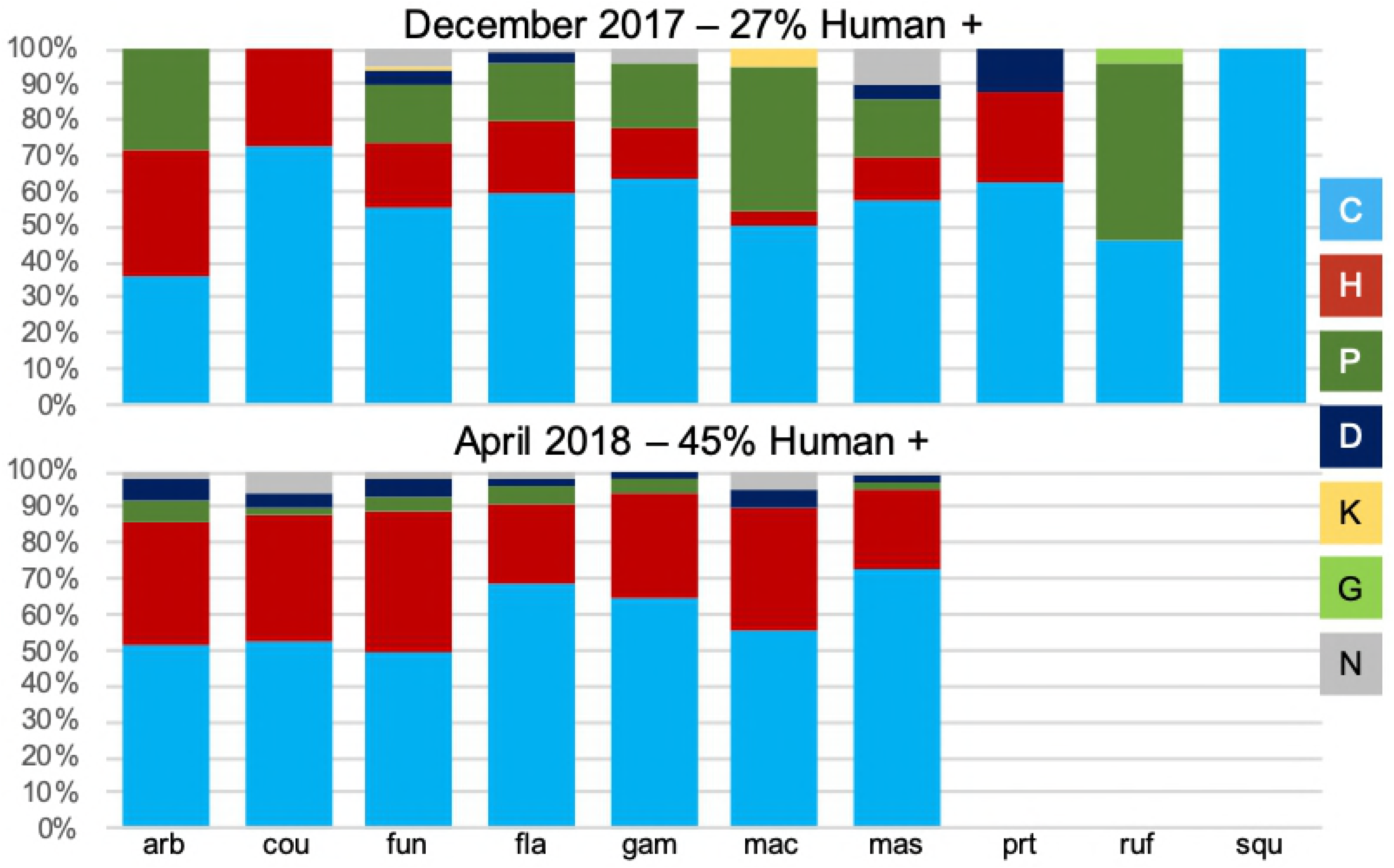
*Anopheles* Host Choice. Proportions represent the overall percentage of positive bloodmeals for respective host species. Therefore, individual bloodmeals with more than one species are counted for each species present. The overall percentage of human positive bloodmeals was 27% in December 2017, and 45% in April 2018. *An. mascarensis:* H = 23, C = 31, P = 7, D = 3, K = 0, G = 0, none = 1; *An. arabiensis:* H = 30, C = 48, P = 1, D = 3, K = 0, G = 0, none = 5; *An. coustani:* H = 226, C = 362, P = 59, D = 31, K = 2, G = 0, none = 24; *An. squamosus:* H = 72, C = 217, P = 37, D = 9, K = 0, G = 0, none = 5; *An. rufipes:* H = 44, C = 139, P = 29, D = 2, K = 0, G = 0, none = 5; *An. funestus:* H = 8, C = 22, P = 9, D = 1, K = 1, G = 0, none = 1; *An. maculipalpis:* H = 68, C = 241, P = 27, D = 11, K = 0, G = 0, none = 16; *An. pretoriensis:* H = 2, C = 5, P = 0, D = 1, K = 0, G = 0, none = 0; *An. gambiae:* H = 0, C = 13, P = 14, D = 0, K = 0, G = 1, none = 0; *An. flavicosta*: H = 0, C = 1, P = 0, D = 0, K = 0, G = 0, none = 0.

The distribution of host bloodmeals identified on the QUEST varied markedly between village and timepoint (Fig 7, S5 Table). The presence of human-positive bloodmeals increased in April 2018 for both villages, from 4% to 48% in Amparihy and 32% to 41% in Ambolodina. Amparihy also exhibited shifts in the number of bovine-positive (96% to 73%) and porcine-positive (96% to 7%) bloodmeals from December 2017 to April 2018. Ambolodina showed a decrease in porcine-positive bloodmeals (22% to 1%), while bovine-positivity increased (88% to 99%). The presence of human-only and pig-only bloodmeals was restricted to traps from Amparihy. Furthermore, human-only bloodmeals were observed across all traps and all quadrants in the April 2018 collection.

**Fig 7.**
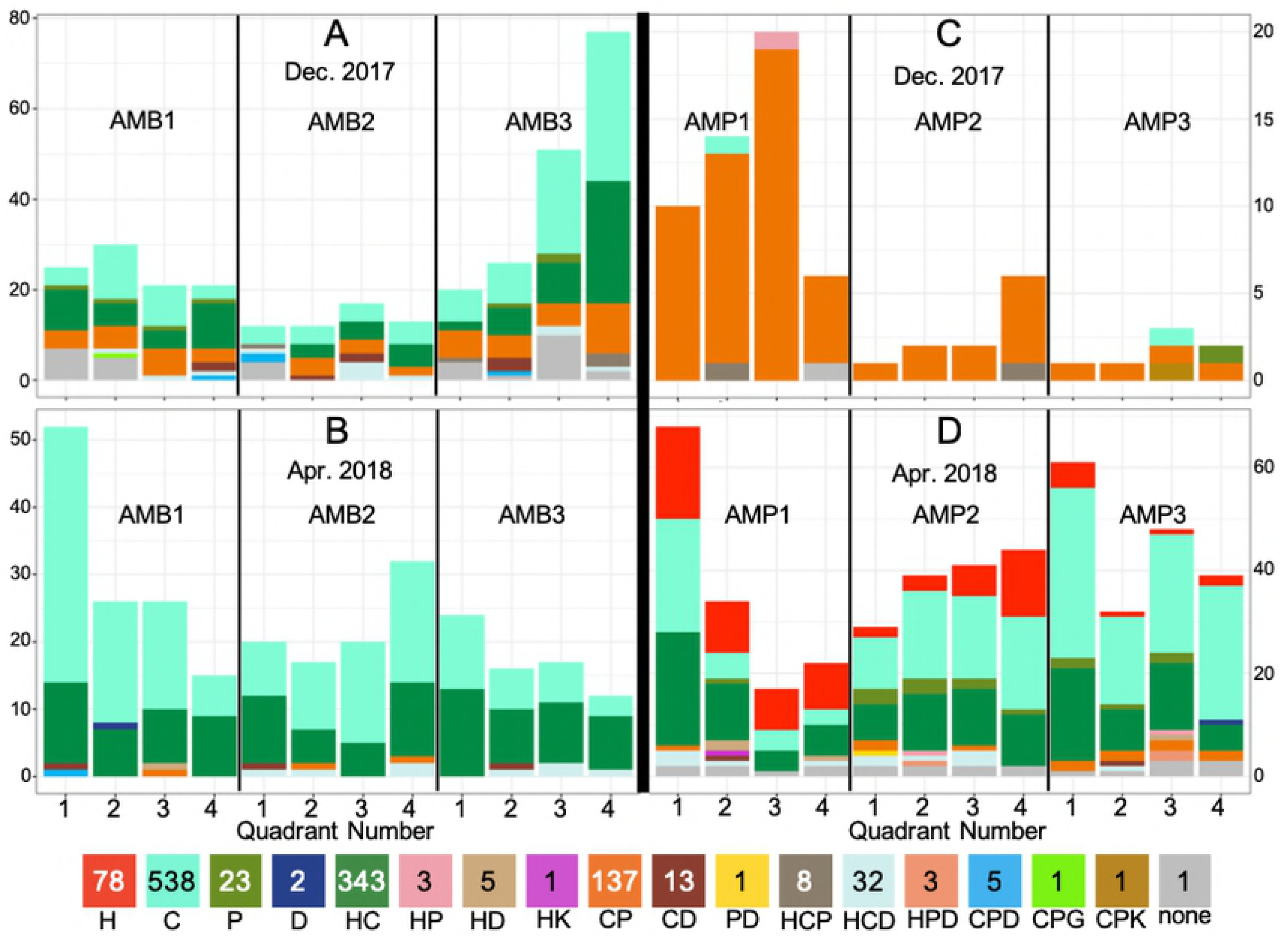
Distribution of hosts across QUEST. H = human, C = cow, P = pig, D = dog, K = cat, G = goat. Letter combinations indicate the presence of two or three hosts in the bloodmeal. Quadrant numbers refer to the direction of their orientation: 1-Northwest, 2-Northeast, 3-Southwest, 4-Southeast. (A) Ambolodina December 2017, (B) Ambolodina April 2018, (C) Amparihy December 2017, (D) Amparihy April 2018. Numerical version available in S5 Table.

Because of the sudden appearance of human-only bloodmeals in Amparihy in April 2018, we wanted to test if there was any morphological difference between mosquitoes classified as having exclusively human-positive bloodmeals and mosquitoes with exclusively cow-positive bloodmeals. Prior studies, focused on Tabanid flies, demonstrated a correlation between mean proboscis length and the mean hair thickness of the biting site [40]. Therefore, we tested whether proboscis length in *An. coustani* was associated with being positive for human-only (n = 58, thinner hide) versus positive for cow-only (n = 59, thicker hide). While there was a slight shift toward a smaller proboscis in human-only mosquitoes, there was no significant relationship between proboscis length and host choice (Fig 8).

**Fig 8.**
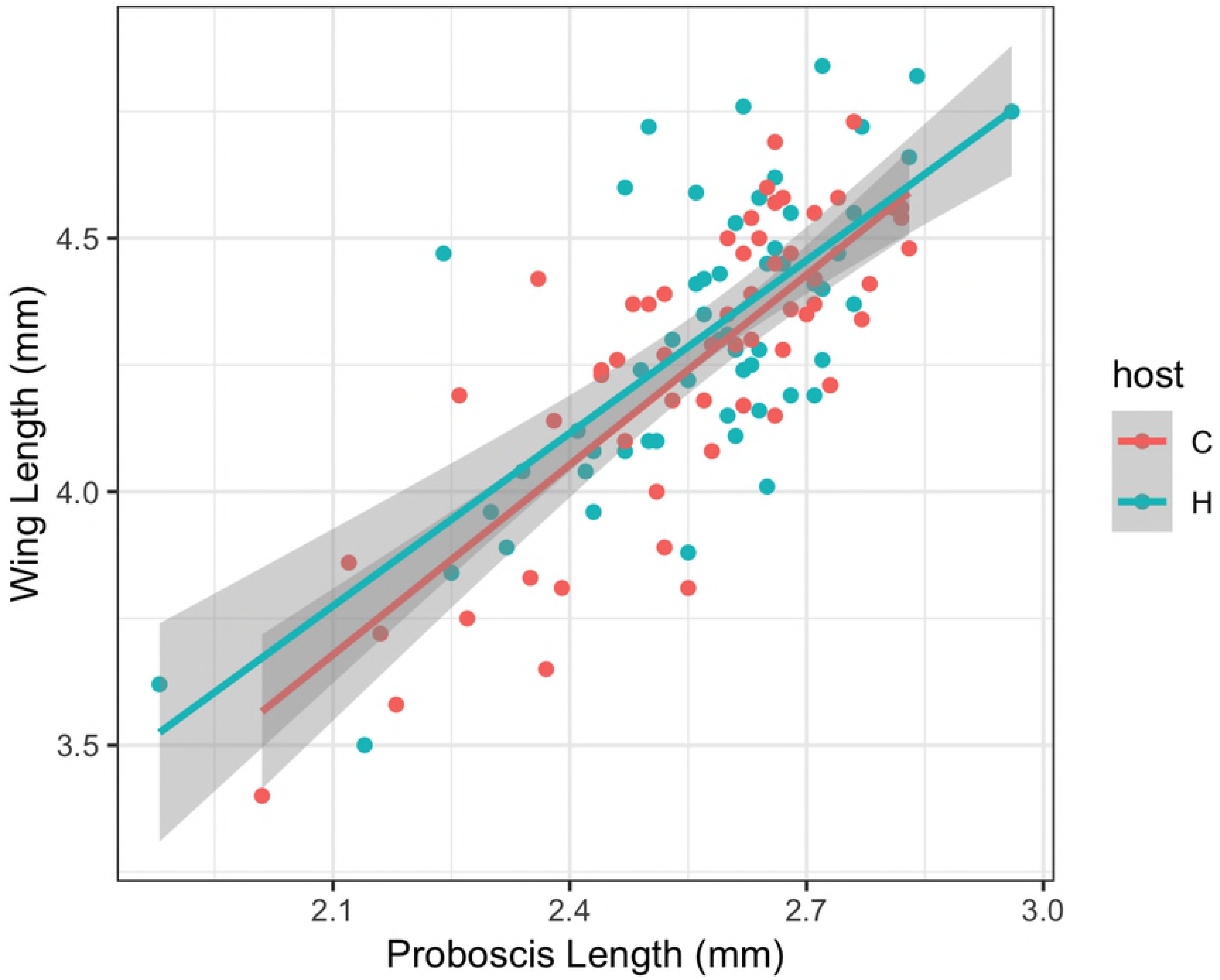
Scatter plot of proboscis length vs. wing length. Proboscis and wing length were measured in millimeters for 117 female *Anopheles coustani* mosquitoes (58 human-positive only [H], 59 cow-positive only [C]). Gray shading around the best fit line represents 95% confidence intervals.

### *Plasmodium* species infection

All four common human *Plasmodium* species (*P. falciparum, P. vivax, P. malariae*, and *P. ovale*) were detected in our assay (Fig 9A) with a total of 75 positive among 1252 collected mosquitoes (6%). The predominant parasites were *P. falciparum* (n=50) and *P. vivax* (n=19), followed by *P. malariae* (n=10) and *P. ovale* (n=8). There was a total of 12 mixed *Plasmodium* species infections, including the combinations *P. falciparum*/*P. vivax* (n=9), *P. falciparum*/*P. malariae* (n=1), *P. falciparum/P. ovale* (n=1), and *P. malariae/P. ovale* (n=1). *Plasmodium*-infected bloodmeals were commonly observed across *Anopheles* species that the Madagascar NMCP typically consider to be secondary or non-vectors, such as *An. coustani* (n=26; 6% of screened *An. coustani* specimens), *An. maculipalpis* (n=20; 8%), *An. squamosus* (n=13; 7%), and *An. rufipes* (n=8; 6%). This constitutes infection rates comparable to, or greater than, rates in the primary malaria vectors in the same sample: *An. funestus* (n=1; 4%), *An. arabiensis* (n=4; 6%), and *An. gambiae* (n=0; 0%). The December 2017 collection contained 18 of the 19 overall observed *P. vivax* infected mosquitoes. The abundance of positive samples, and diversity of *Plasmodium* spp. within them, shifted in these two villages, with *P. falciparum* comprising nearly all of the *Plasmodium* positive bloodmeals in April 2018. The village of Amparihy had no *Plasmodium-positive* mosquitoes in December 2017, while similar numbers of Plasmodium-positive mosquitoes were observed between the villages in April 2018 (Fig 9B). Apart from this difference in *Plasmodium* positive mosquito distributions between Amparihy and Ambolodina, no specific pattern of *Plasmodium* positive mosquitoes was observed across QUESTs (S1 Fig, S6 Table).

**Fig 9.**
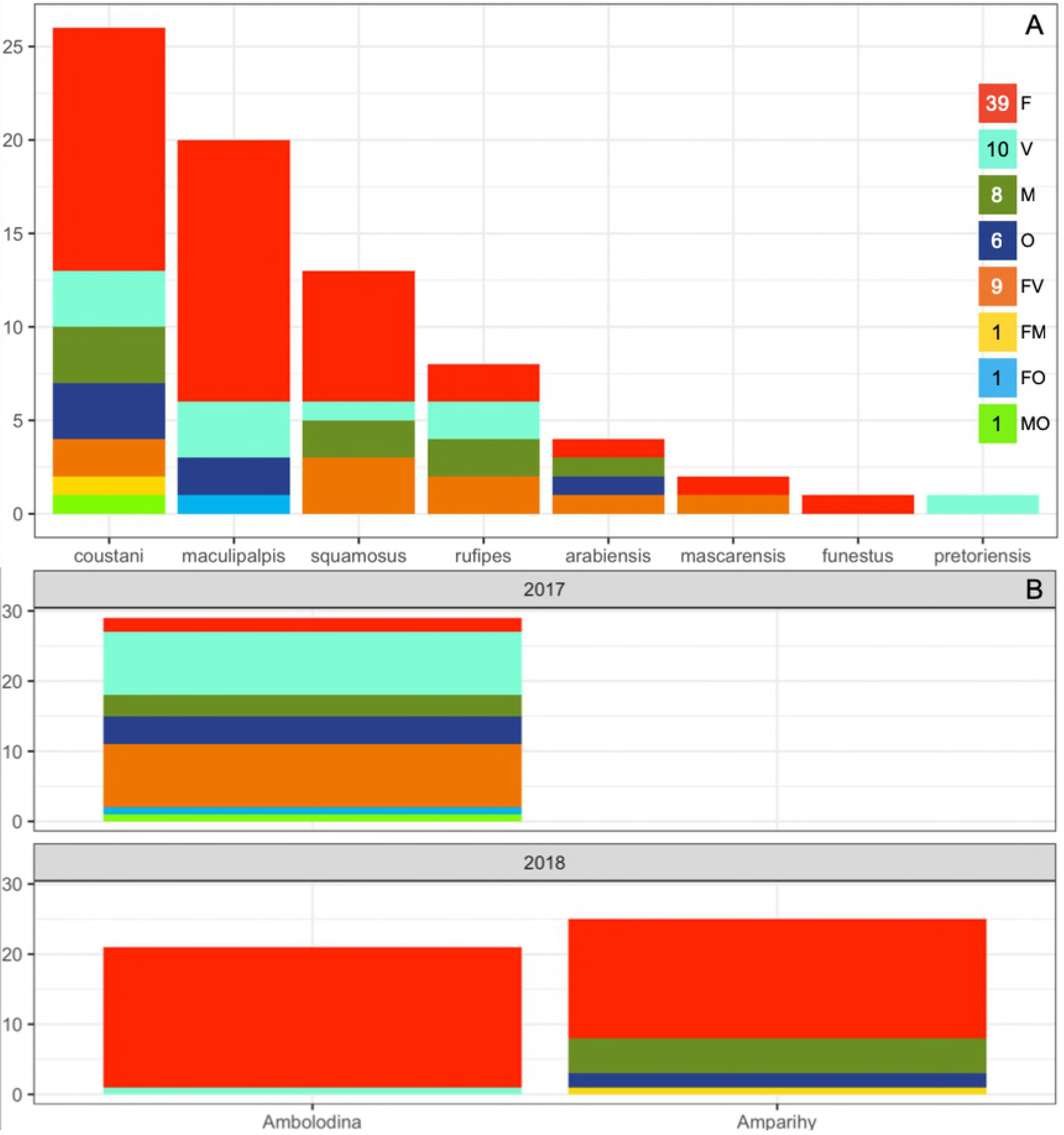
Distribution of *Plasmodium* positive mosquitoes. F =*P. falciparum*, V =*P. vivax*, M =*P. malariae*, O =*P. ovale*. Letter combinations indicate the presence of multiple *Plasmodium* species in the bloodmeal. (A) *Plasmodium* species infection across *Anopheles* species. (B) *Plasmodium* positive mosquitoes across the villages of Ambolodina and Amparihy in December 2017 and April 2018.

## Discussion

This study demonstrates the ability of QUESTs and BLOODART to refine perspectives on mosquito collection and assessment. QUESTs collect a diverse and substantial sample of *Anopheles* mosquitoes with a simple design. Many of the components can be sourced directly from the environment for the initial setup or repair. By collecting additional data from the surrounding environment (such as a host census including humans and domesticated animals, weather conditions, and nearby breeding sites), we could obtain further insight into the behavior of these medically important mosquitoes. The data produced by a BLOODART analysis of captured mosquitoes augments the utility of QUESTs (or any other mosquito trap strategy), efficiently monitoring additional important vector indicators. By analyzing all of our captured Anopheline mosquitoes, as opposed to focusing only on the predetermined important vectors, we detected a significant parasite reservoir in the abdomens of secondary vectors. In the context of limited time and resources in outbreak or surveillance scenarios, secondary vectors may be passed over for assessment of vector indicators, or never collected at all. This highlights the problematic nomenclature surrounding “primary” or “secondary” vectors, in that they may constrain a thorough investigation of complex disease transmission networks. The collection and analytical tools used here make comparisons across mosquito species feasible.

As a result of this study, and training missions carried out at the Madagascar NMCP, the skillset necessary to use both QUEST and BLOODART is currently in place. This study provides a framework for how these tools can be deployed to enhance Madagascar’s capacity for entomological surveillance and malaria elimination.

### Mosquito Capture

QUEST collection was effective at capturing a diverse sample of female blood fed *Anopheles* mosquitoes. Placing the QUESTs in the same locations in December 2017 and April 2018 allowed for comparison of QUEST data across the two timepoints. The QUESTs AMB1, AMB3, and AMP1 had significantly different mosquito distributions across the four quadrants between the December 2017 and April 2018 timepoints. In December 2017, the traps AMB3 and AMB4 had non-homogenous mosquito distributions across the QUEST quadrants. In April 2018, the traps AMB1 and AMP1 had non-homogenous mosquito distributions across the QUEST quadrants (Fig 2, S2 Table). This study did not perform a formal comparison between QUEST and standard collection methods (such as human landing catch or CDC light traps), or a direct comparison to the standard barrier screen or pyrethroid spray catch, but proved to be far higher-yielding and less obtrusive than the latter. The mosquito diversity of the indoor pyrethroid spray catch was limited to a single species, *An. funestus*,while the QUEST captured this species in addition to nine others. The standard barrier screen captured a substantial number of mosquitoes per sampling night, likely due to its placement directly inside a Malagasy zebu corral with a high concentration of available bovine hosts. While the QUESTs often were not far from where cattle or pigs were corralled, they were never placed directly inside them, granting insight into a potentially less biased distribution of species and hosts. The standard barrier screen captured seven of the 10 species captured on the QUESTs. The species that were not captured on the standard barrier screen had low capture numbers overall (S3 Table), suggesting that these species may have surfaced given a greater number of sampling nights.

### Mosquito Characterization

Concordance between morphological and BLOODART identifications ranged from 93-99%. Unique specimen vouchers enabled us to refer back to discordant specimens, which revealed that morphological misidentification was responsible for many of the discrepancies between morphological and molecular identifications. Field conditions (such as poor lighting) likely contributed to less than optimal species identifications. Additionally, vouchered specimens allowed us to associate a novel *Anopheles* ITS2 sequence probe [25] with the morphologically identified species *An. mascarensis*, facilitating the first online deposition of genetic data for this common Malagasy species (Genbank accession: MH560267). By vouchering, photographing, and depositing our mosquito data on VectorBase, we have secured the future utility of our mosquito collection for the purpose of further morphological and molecular investigations.

### Mosquito Abundance and Diversity

The predominant mosquito species in this collection have been considered vectors of secondary importance. However, their abundance relative to primary vectors necessitates closer examination of their potential impact on human health. *Anopheles coustani*, comprising 37% of our mosquito collection, was recently implicated as a vector of malaria [10]. Despite being primarily exophagic and exophilic, overwhelming numbers could result in this species being responsible for the majority of indoor bites despite the presence of endophagic and endophilic species [10]. Similarly, *An. maculipalpis, An. rufipes, An. squamosus*, and *An. mascarensis* have all been documented with *Plasmodium* parasites [16–22]. The relative abundance of these species in conjunction with their malaria transmission potential warrants further investigation.

*Anopheles gambiae s.s*. is uncommon in the highlands of Madagascar [42], with *An. arabiensis* being the predominant representative of the *An. gambiae s.l*. complex in this region. Permanent shifts in species dominance from *An. gambiae s.s*. to *An. arabiensis* have been reported in continental Africa [43,44]. Typically, this is viewed as a succession event, a product of the differential effects of indoor residual spraying and insecticide treated nets on these two species. These mosquito control strategies are most effective against the endophilic/endophagic *An. gambiae s.s*., with negligible impacts on the exophilic/exophagic *An. arabiensis* [43,44]. The habitat preferences for these two species differ as well, with *An. gambiae s.s*. preferring a more humid climate, while *An. arabiensis* is found in both humid and arid environments [5,19,45,46]. Considering that our timepoints occur at the very beginning and end of the 2017-2018 rainy season (considered to be the malaria transmission season in Madagascar [34]), we might naturally expect to see seasonal variation in species occurrence and abundance. Further, longitudinal monitoring of these sites would likely provide more insight on the population dynamics of these closely related species. Insecticide treated nets were observed to be virtually absent from these villages. Consequently, it was not surprising that the mosquitoes collected here did not exhibit the behavioral adaptation of feeding earlier in the evening [47].

There are several phenomena that limit direct comparison of mosquito data between villages. Due to logistical issues ranging from inclement weather to security, the number of nights sampled in each village differed. Further, villages were sampled consecutively as opposed to concurrently, introducing a unique mixture of irretrievable ecological factors into each trapping night across the overall sampling period.

The distribution of mosquitoes collected from the QUEST is potentially influenced by wind direction, precipitation, temperature, lunar cycle, proximity to breeding sites, and distance to viable hosts [32,48–50]. Specific data on these variables were not collected, limiting our conclusions. Numerous mosquitoes were caught in the eaves of the trap. Mosquitoes tend to move up and over physical barriers [32], suggesting that the eaves may play a role in prolonging mosquito detainment or improved capture. The height at which mosquitoes were captured was not recorded.

### Host Choice

The mosquitoes in this survey exhibited complex feeding behaviors. Individual mosquito species frequently fed on human and non-human hosts, even among species considered highly anthropophilic. As demonstrated in a metanalysis of the human blood index in *An. gambiae* sibling species, sampling site (indoors vs outdoors) is more closely associated with human positive bloodmeals than mosquito species [51]. The higher diversity of host choice in this study may reflect a reduction in sampling bias for endophilic vector species like *An. gambiae s.s*. and *An. funestus*. The BLOODART analysis revealed multiple blood feeding behavior (46.4%) in our mosquito sample. PCR has the capacity to detect blood consumed 36-48 hrs after ingestion [25,52,53], lasting through the feeding phase of the gonotrophic cycle [54]. The increased incidence of multiple blood feeding, as compared to most surveys, could be a difference in bloodmeal assessment technique, with many studies choosing to pursue a narrower range of possible hosts [26,30,55,56]. High incidence of multiple blood feeding has important epidemiological implications. Numerous mosquitoes showed evidence of feeding on humans and one or two additional mammal species. This may increase the probability that a mosquito is also feeding on multiple individuals within a species, amplifying the vectorial capacity of the potential malaria vectors in these villages. The zoophilic preferences of the mosquitoes in this sample, however, may temper their impact on malaria transmission by dedicating potentially infective bites to non-human hosts. There was a substantial shift toward human feeding in the April 2018 collection, primarily observed in the species *An. coustani*, *An. funestus*, *An. rufipes*, and *An. arabiensis*. An increase in human feeding could lead to an increase in the presence of *Plasmodium* positive mosquitoes, though evidence of such from this study was variable between villages (Fig 9B). The possible reasons for this anthropophilic shift include host availability [41,57,58], insecticide treated net coverage [59,60], or perhaps a plastic trait influencing host choice. Based on prior studies demonstrating a correlation between mean proboscis length and the mean hair thickness of the biting site chosen in Tabanid flies [40], we tested the idea that proboscis length may be a plastic trait influencing host choice. In a comparison of human-only fed and cow-only fed *An. coustani*, there was a non-significant shift toward a smaller proboscis in the human fed mosquitoes. Our sample, however, was skewed toward larger proboscis mosquitoes. If we had a more representative sample of larger and smaller proboscis lengths, we may have detected a significant difference in host choice.

At times, there may be approximately as many Malagasy zebu as humans in these villages, characteristic of much of the region [45]. Mosquitoes will target hosts with a greater surface area or weight [61], and considering the relative size of the Malagasy zebu, the 90.3% prevalence of bovine blood may be expected. Frequent feeding between non-human hosts, both within and between species, could increase the risk for outbreaks of mosquito-borne epizootic diseases. There were numerous pigs, semi-domesticated dogs, and cats present in both villages. Goats were spotted several kilometers from the villages, but none were observed within them. Lemurs and rodents were not observed.

The distribution of host choice across mosquitoes on the nets (Fig 6) may be influenced by the proximity of hosts to the net. It is also likely influenced by the composition of mosquito species on individual QUESTs, each of which displayed an individual hierarchy of host preferences (Fig 7). Although no empirical census of hosts was conducted, there did anecdotally appear to be more pigs in Amparihy and more cattle in Ambolodina, potentially explaining the higher percentage of pig and cattle DNA, respectively, in the abdomens of mosquitoes from these villages.

### *Plasmodium* species infection

The incidence of *Plasmodium* infection for the mosquitoes collected in this study (6%, n=75) likely reflects our unbiased approach to mosquito collection and processing, as we did not limit our molecular analysis to predetermined primary vector species. However, we should consider that *Plasmodium* positivity in this assay is observed in the abdomen of the mosquito, and does not indicate the status of sporozoites in the mosquito’s salivary glands. An infectious mosquito is typically defined as a mosquito with sporozoites in the head/thorax [62], which, due to daily rates of mortality [63], will only be a portion of the mosquitoes with the earlier parasite stages in the gut [64]. All of the mosquito species in our sample have been previously documented with sporozoites in Madagascar [10,14,16–22], and the predominant species observed here, *An. coustani*, was recently implicated as an important vector of *Plasmodium* parasites [10] in the region. The combined impact of *An. coustani* and the remaining secondary vectors may be responsible for sustained malaria transmission in these villages, which have low primary vector density. Consequently, we have identified a potential reservoir of *Plasmodium* parasites outside the realm of many current vector interventions. Therefore, targeted efforts focused on the suppression of *An. funestus* and *An. gambiae s.l*., which primarily include indoor interventions, may not be sufficient to effectively reduce malaria transmission.

We revealed differences in the occurrence of *Plasmodium* spp. positivity across space and time. This may reflect the seasonality of *P. falciparum* and *P. vivax* malaria, which peaks in the study region around the time of the second round of surveys described here [34]. Our mosquito data reflect a pattern of *P. vivax* positivity restricted to the beginning of the rainy season, and a predominance of *P. falciparum* toward the end of the rainy season. The distribution of *Plasmodium* positive mosquitoes on the nets was fairly homogeneous, with the exception of the village of Amparihy in December 2017, which had no positive mosquitoes. The observation of a statistically significant pattern is limited by the total number of positive mosquitoes (n=75) distributed across 48 possible quadrants.

### Conclusion

To achieve the goal of malaria elimination in this country, the role of neglected secondary vector species must be considered. The lack of comprehensive surveillance, artemisinin combination therapy drugs, insecticide treated nets, and rapid diagnostic tests in remote areas of the country provide a safe refuge for *Plasmodium* parasites. Substantial parasite reservoirs in neglected vector species adds to the series of gaps that expose hard-earned malaria-free districts to the perpetual threat of recrudescence.

## Declarations

Ethics approval and consent to participate

## Acknowledgements

The authors thank the community of the study villages for their willingness to support our work, to the ASA (Ankohonana Sahirana Arenina, www.asa-madagascar.org) organization for permission to conduct our surveys, to PACT Madagascar (www.pact-madagascar.org) for logistical support, and to the DLP (Direction de Lutte contre le Paludisme) for support in carrying out field operations. We thank Jean-Claude, Heriniana, Mirar, and Teddy for their assistance in conducting the field collections with RET. Many thanks to Dr. Rajeev Mehlotra, Dr. Rosalind Howes, Estee Cramer, Brooke Roeper, Marlin Linger, and the Ecology and Evolution Journal Club at CWRU for helpful comments on early drafts of the manuscript. We also thank Iulian Gherghal for suggesting that proboscis length may influence host choice, inspiring the authors to pursue this question.

## Supporting Information

**S1 Fig) Distribution of *Plasmodium* positive mosquitoes across QUEST in December 2017 and April 2018**. F = *P. falciparum*, V = *P. vivax*, M = *P. malariae*, O = *P. ovale*. Letter combinations indicate the presence of two or three species in the bloodmeal. Quadrant numbers refer to the direction of their orientation: 1-Northwest, 2-Northeast, 3-Southwest, 4-Southeast. (A) Ambolodina December 2017, (B) Ambolodina April 2018, (C) Amparihy December 2017, (D) Amparihy April 2018. Numerical version available in S6 Table.

**S1 Table) BLOODART Ligase Detection Reaction probes for the detection of mammalian host and mosquito species.**

**S2 Table) Mosquito abundance across QUEST.**

**S3 Table) Comparison of capture numbers between QUESTs and standard barrier screen.**

**S4 Table) Distribution of mosquito species across QUEST.**

**S5 Table) Distribution of mosquito hosts across QUEST.**

**S6 Table) Distribution of mosquito Plasmodium infection across QUEST.**

## Data Reporting

VectorBase (VBP0000345)

### Accession Numbers

Genbank accession for *Anopheles mascarensis:* MH560267

### Striking Image

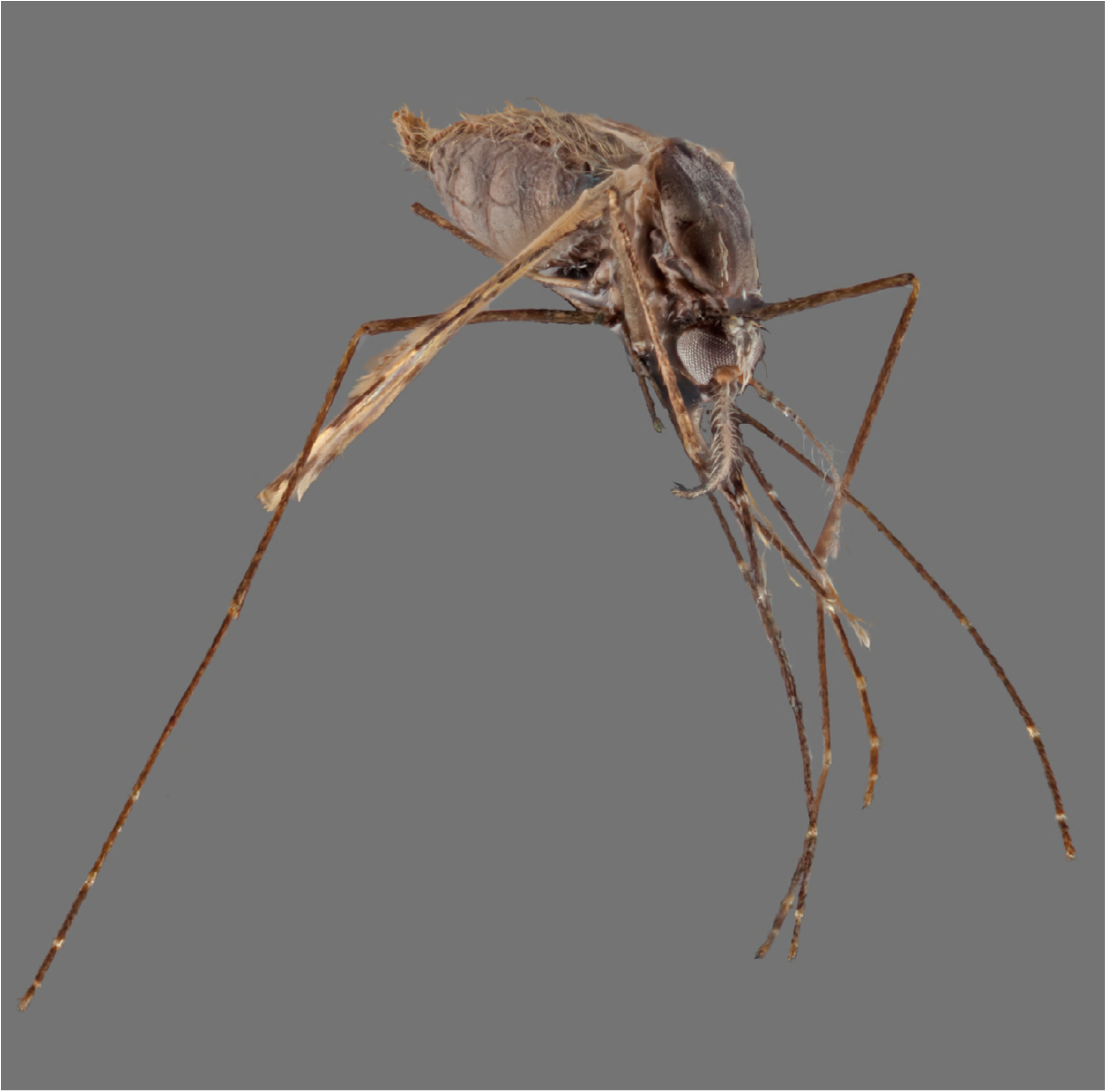

## Additional Info for Submission

### Financial Disclosure

Stipend support for R. E. Tedrow was provided by the United States Navy Health Services Collegiate Program. Additional support was provided to P. A. Zimmerman through the CWRU School of Medicine.

### Related Manuscripts

### Reviewer/Editor Suggestions (4)

**Tom Burkot; Professor**

James Cook University; Australian Institute of Tropical Health and Medicine

**Neil Lobo; Research Associate Professor**

University of Notre Dame; Department of Biological Sciences

**Laura Harrington; Professor**

Cornell University; Department of Entomology

**Kate Altman; Director of Research**

UT Health San Antonio; School of Health Professions

**Authors’ contributions**
RET, TN, JR, ACR, GJS, and PAZ participated in study and trap design. RET, TR, and TN carried out field activities in Madagascar. RET and PAZ designed the assay and RET processed the samples and analyzed the data. RET produced the first draft of the manuscript. RET, TR, TN, JR, ACR, GJS, and PAZ contributed substantially to revision of the first draft.

